# M2 Macrophages are Major Mediators of Germline Risk of Endometriosis and Explain Pleiotropy with Comorbid Traits

**DOI:** 10.1101/2024.11.21.624726

**Authors:** Soledad Ochoa, Fernanda S. Rasquel-Oliveira, Brett McKinnon, Marcela Haro, Sugarniya Subramaniam, Pak Yu, Simon Coetzee, Michael S. Anglesio, Kelly N. Wright, Raanan Meyer, Caroline E. Gargett, Sally Mortlock, Grant W. Montgomery, Michael S. Rogers, Kate Lawrenson

**Affiliations:** Department of Obstetrics and Gynecology, University of Texas Health, San Antonio TX, USA; Center for Inherited Oncogenesis, University of Texas Health, San Antonio TX, USA; Vascular Biology Program, Boston Children’s Hospital, Department of Surgery, Harvard Medical School, Boston, MA, USA; The Institute for Molecular Bioscience, The University of Queensland, Brisbane, Australia; Women’s Cancer Research Program at the Samuel Oschin Comprehensive Cancer Center, Cedars-Sinai Medical Center, Los Angeles, CA, USA; British Columbia’s Gynecological Cancer Research (OVCARE) Program, University of British Columbia, Vancouver General Hospital, and BC Cancer, Vancouver, British Columbia, Canada; Department of Obstetrics and Gynecology, UBC, Vancouver, British Columbia, Canada; Division of Minimally Invasive Gynecologic Surgery, Department of Obstetrics and Gynecology, Cedars-Sinai Medical Center, Los Angeles, CA, USA; The Ritchie Center, Hudson Institute for Medical Research, Melbourne, Victoria, Australia; Department of Obstetrics and Gynaecology, Monash University, Melbourne, Victoria, Australia

**Keywords:** Endometriosis, GWAS, inflammation, M2 macrophages, IL1A, IL1B, single cell transcriptomics

## Abstract

Endometriosis is a common gynecologic condition that causes chronic life-altering symptoms including pain, infertility, and elevated cancer risk. There is an urgent need for new non-hormonal targeted therapeutics to treat endometriosis, but until very recently, the cellular and molecular signatures of endometriotic lesions were undefined, severely hindering the development of clinical advances. Integrating inherited risk data from analyses of >450,000 individuals with ∼350,000 single cell transcriptomes from 21 patients, we uncover M2-macrophages as candidate drivers of disease susceptibility, and nominate IL1 signaling as a central hub impacted by germline genetic variation associated with endometriosis. Extensive functional follow-up confirmed these associations and revealed a pleiotropic role for this pathway in endometriosis. Population-scale expression quantitative trail locus analysis demonstrated that genetic variation controlling *IL1A* expression is also associated with endometriosis risk variants. Manipulation of IL1 signaling in state-of-the-art *in vitro* decidualized assembloids impacted epithelial differentiation, and in an *in vivo* endometriosis model, treatment with anakinra (an interleukin-1 receptor antagonist) resulted in a significant, dose-dependent reduction in both spontaneous pain and evoked pain. Together these studies highlight non-diagnostic cell types as central to endometriosis susceptibility and support IL1 signaling as an important actionable pathway for this disease.

## Introduction

Endometriosis is a common yet poorly understood gynecological disease, affecting around 10% of biologically female individuals at reproductive-age. In endometriosis, glands containing endometrial-like epithelium and stroma are found outside of the uterine cavity, often within the ovaries and throughout the peritoneal cavity.

Endometriosis causes chronic pain, dysmenorrhea and infertility, and is associated with an increased risk of rare types of ovarian cancer ^4,5^. While the presence of endometrial-type epithelium and stroma are diagnostic for disease, other microenvironmental cell types play essential roles in disease pathogenesis. Fibrosis and dysregulated innate and adaptive immune responses are other hallmarks of disease ^6,7^. Recent single cell transcriptomic studies have profiled diagnostic and microenvironmental components of endometriosis^8,9^ yielding new insight into the cell types, cell states, genes and pathways involved in endometriosis pathophysiology ^1–3^.

Genome-wide association studies (GWAS) profile common genetic polymorphisms across population-scale cohorts to identify genetic variants associated with susceptibility to complex traits. GWAS have identified thousands of associations that impact risk for thousands of phenotypes, including endometriosis ^10–13^. The era of GWAS spawned a large field of ‘post-GWAS’ functional projects aiming to construct the pathways by which risk variants, typically located in noncoding DNA, perturb gene expression to impact disease susceptibility ^14^. For endometriosis risk variants, a number of studies have quantified mRNA expression, miRNA expression or methylation in eutopic endometrium to identify candidate genes, particularly those expressed in epithelial cells, that are associated with susceptibility to endometriosis ^15–18^. Here we extend this work by integrating single cell landscapes of endometriosis to explore the role of the microenvironment in endometriosis risk. We report a central role for myeloid cells in endometriosis risk that may also explain genetic overlaps with certain cancers, pain conditions and inflammatory traits.

## Results

### Single cell disease risk scores associate M2 macrophages and endometriosis risk

Given the known importance of immunity and inflammation in endometriosis, we annotated immune subsets in reanalyzed data from a recent single cell atlas of peritoneal endometriosis (n=32 samples), endometrioma (n=8 samples), endometriosis-free ovaries (n=4 samples) and eutopic endometrium (n=10 samples) from 21 patients ^8^. A total of 118,103 CD45-positive (*PTPRC*-expressing) immune cells were stratified into 15 functionally distinct populations (see Methods and Supplemental Table 1). T cell subsets included 5 clusters of CD8 cells (63,906 cells), gamma-delta T cells (γδ; 16,284 cells), Th2 CD4 T-cells (6,843 cells) and Th17 CD4 T-cells (920 cells). Myeloid subsets included 8,790 dendritic cells, 1,139 M1 macrophages and 5,554 M2 macrophages. The remaining subsets were B cells (5,274 cells), plasmablast/plasma cells (1,771 cells), natural killer (NK) cells (5,977 cells), and mast cells 1,645 cells ^8,9^. Other cell types represented in the data set include the endometrial-type epithelium (8,160 cells) and endometrial-type stroma (66,721 cells) that comprise the glandular structures seen in eutopic endometrium and endometriosis, plus mesenchymal cells (125,311 cells), mesothelial cells (2,865 cells), and endothelial cells (23,226 cells) (Fig. 1A-C).

**Fig. 1:**
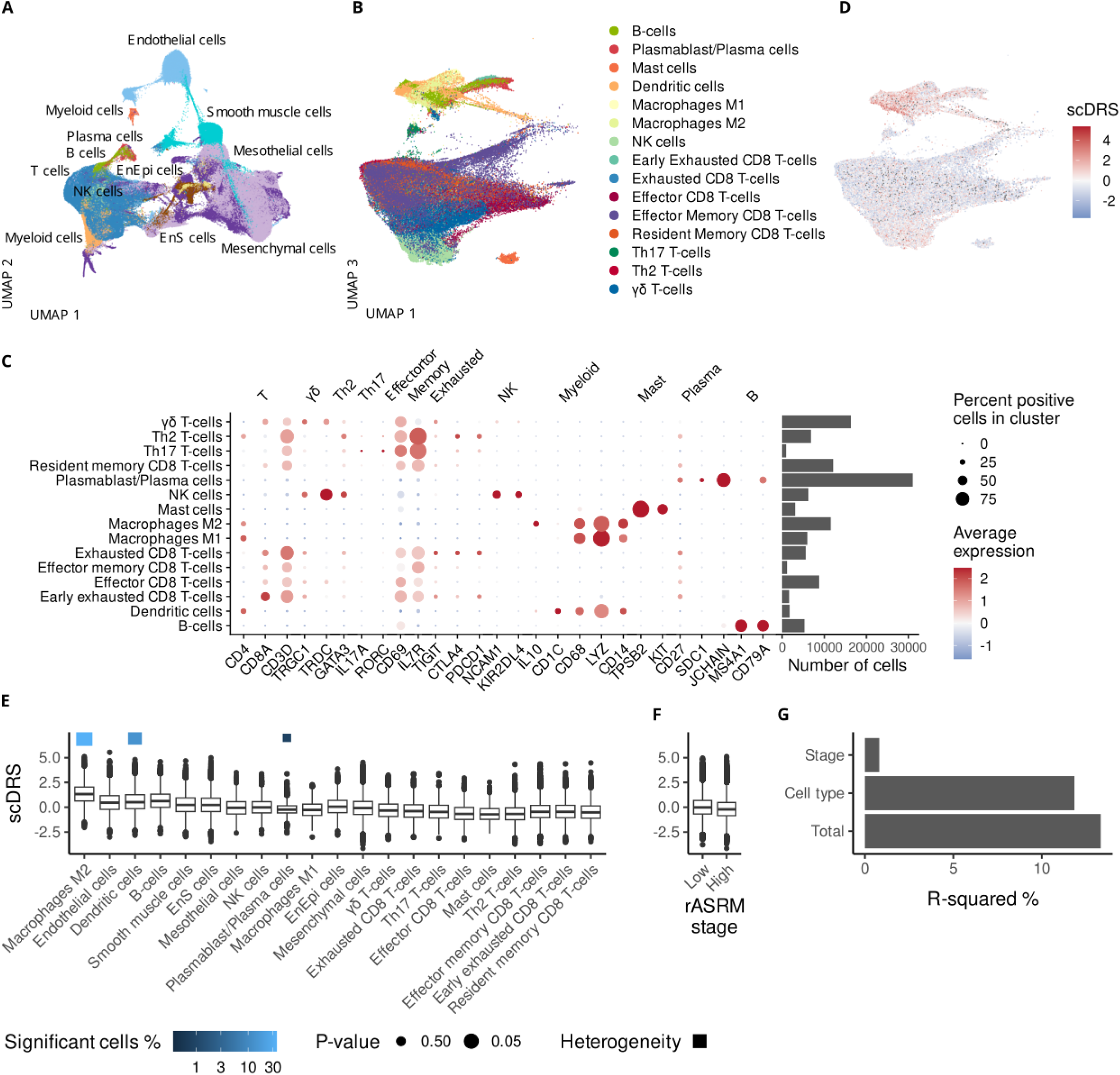
Endometriosis risk gene expression is enriched in M2 macrophages, dendritic cells and plasmablasts/plasma cells. A) UMAP representation of the major cell types. The data set includes 362,700 cells from 49 samples taken from 21 patients. Erythrocytes are excluded. B) UMAP representation of the immune subclusters present. C) Canonical markers identifying the immune clusters. (D-F) Distribution of the single cell disease relevance score in D) the UMAP space, E) per cell cluster, F) per Stage. Only the 200,798 cells from endometrioma and peritoneal endometriosis are included here. G) scDRS variance explained by different cell categories according to linear regression. UMAP, uniform manifold approximation and projection; NK, natural killer cells. In E) and F) upper and lower bounds of boxes represent upper and lower quartiles, with central lines at the median. Solid lines above and below the boxes denote the minimum and maximum data points, and black dots denote outliers. Blue squares highlight cell populations with significant heterogeneity in risk scores (*P* < 0.05).

The single cell disease relevance score (scDRS)^1^ algorithm was implemented to identify cell types associated with endometriosis risk by integrating the single cell transcriptome profiles with genome-wide association data from a meta-analysis of 450,668 controls and 23,492 endometriosis cases ^10^. This association study included patients with both minimal/mild (revised American Society for Reproductive Medicine (rASRM) stage I/II) and severe/extensive(revised American Society for Reproductive Medicine (rASRM) stage III/IV); most subjects were of European ancestry. ScDRS identified a significant association with inherited risk of endometriosis and M2 macrophages (Monte Carlo test *P* = 0.013), dendritic cells (Monte Carlo test *P* = 0.050), and endothelial cells (Monte Carlo test *P* = 0.044) (Fig. 1D,E, Supplementary Table 2). B cells did not quite reach statistical significance (Monte Carlo test *P* = 0.051). Within cell-type heterogeneity was detected for M2 macrophages (Monte Carlo test *P* = 0.031), dendritic cells (Monte Carlo test *P* = 0.029), and plasma cells/plasmablasts (Monte Carlo test *P* = 0.045), indicating subsets of these cells drive association with disease (Supplemental Table 2). Although higher stage endometriosis has greater heritability ^19^, when stratifying cells based on stages, we did not see a difference in scDRS score for cells isolated from patients with stage I/II compared to stage III/IV disease (Fig. 1F). Each stage group was represented by 7 patients. Cell type explained 14.65 times more variance in score compared to stage (Fig. 1G).

### Macrophage-derived *IL1A/ILB* as likely mediators of risk conferred by genetic variants at 2q41.1

To better understand the association with M2 macrophages, dendritic cells and endothelial cells, we identified the genes whose expression is correlated with the risk score (Fig. 2A), and the pathways they affect (Fig. 2B). Correlation is centered around 0 for the 30,354 features in the dataset, but 1,642, 3,365 and 4,157 genes are significantly correlated (adjusted *P* < 0.05) with the scDRS score in M2 macrophages, dendritic cells and endothelial cells, respectively. *IL1A* and *IL1B* are the top ranked genes with the highest positive correlation in both myeloid subgroups (*IL1A* spearman correlation 0.59 for M2 macrophages, 0.36 for dendritic cells; *IL1B* correlation 0.51 for M2 macrophages and 0.46 for dendritic cells, adjusted P < 2.61x10^-222^ for each test). The top genes for endothelial cells are different but include genes known to be involved in the disease. Variants upstream of *KDR,* encoding VEGFR2, have been associated with disease severity; CALCRL functions as co-receptor of CGRP, which affects lesion size and pain ^20^ and, finally, the LIF receptor seems to contribute to lesion immunomodulation.^21,22^ (Fig. 2A). At the pathway level, signaling by different interleukins is highly enriched among the positively correlated genes for myeloid cells (adjusted *P* < 1.97x10^-08^) (Fig. 2B). Additionally, translation-related pathways are highly represented, especially among negatively correlated genes for endothelial cells.

**Fig. 2:**
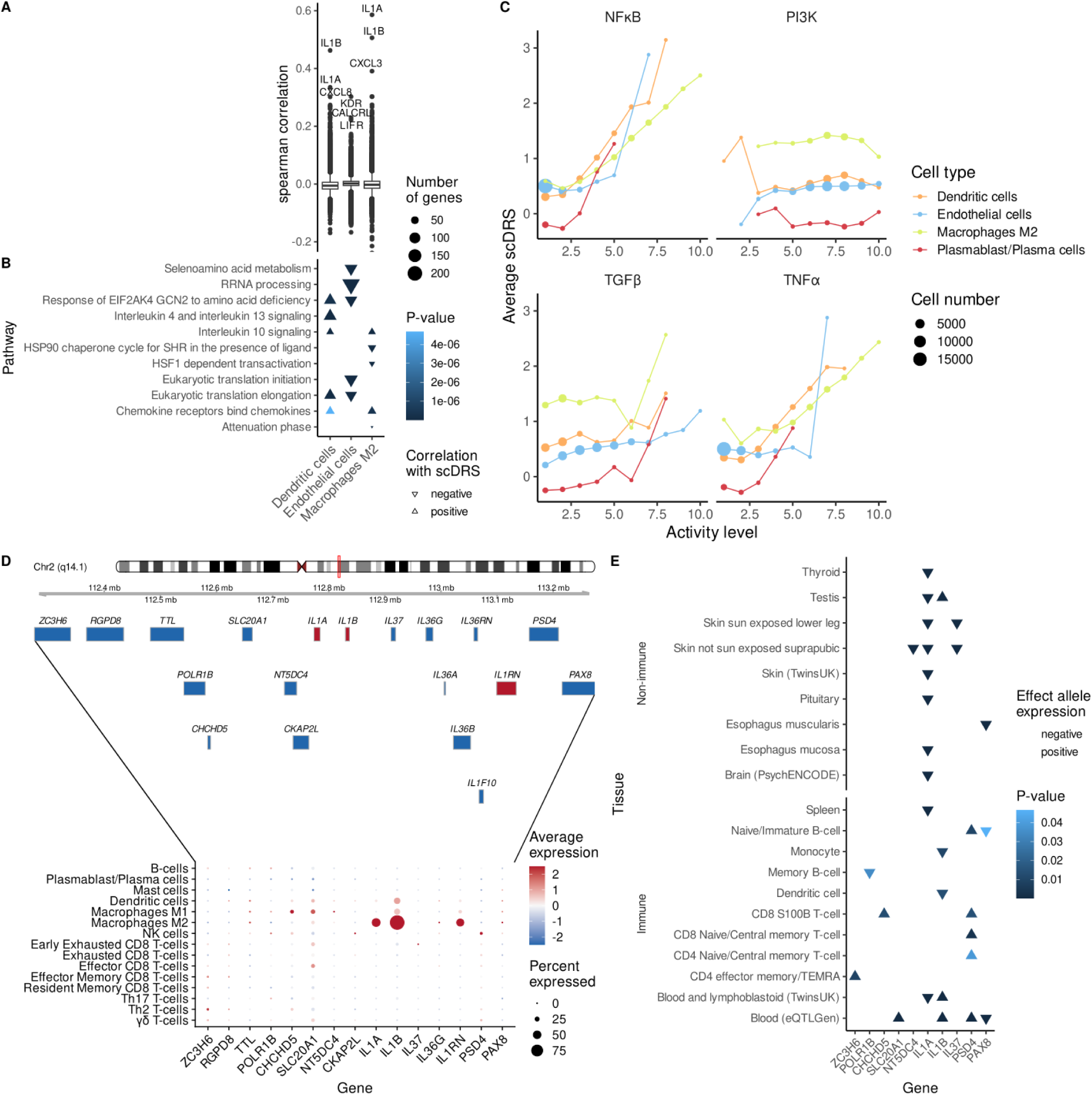
Endometriosis risk is consistently associated with *IL1A/B* gene expression. A) Correlation between gene expression and scDRS. B) Pathways enriched among genes correlated with scDRS. C) scDRS heterogeneity relative to pathway activity. Spearman correlation. D) Gene expression of candidate susceptibility genes at the 2q41.1 endometriosis risk locus in immune subsets. E) Expression quantitative trait locus analysis of differences in candidate risk gene expression associated with endometriosis risk variant genotype.

Of the three cell types associated with endometriosis, both myeloid populations displayed significant within-cell-type disease-association heterogeneity (Fig 1E). To check if this heterogeneity is associated with cell function, we estimated pathway activity using PROGENy scores ^23^. Comparing binned pathway scores against average scDRS, a positive correlation between risk and pathway activity is evident for NF-κB (Spearman correlation = 0.92, *P*-value 2.01x10^-7^) and TNF-α (Spearman correlation = 0.8, *P* 1.2x10^-6^), with M2 macrophages reaching higher levels of activity (Fig. 2C). No other significant correlations between scDRS bin and pathway activity were detected, including for PI3K, a pathway somatically altered in endometriosis ^24^ (Fig. 2c and Supplementary Fig. 1). IL1A and IL1B function in a feedback loop with NFκB signaling ^25^ and have been proposed as the likely causal risk genes at chromosome 2q14.1, a locus associated with endometriosis across multiple populations ^26–28^. Endometriosis risk variants lie around 2kb centromeric to the promoter of *interleukin 1A*. While the locus harbors over a dozen protein coding genes, most candidate risk genes were not expressed in any of the cell types present in the single cell endometriosis atlas (Fig. 2D). *IL1A* and *IL1B* are notable for high expression in M2 macrophages. Endometriosis-associated M2 macrophages also express IL1A/B antagonist *IL1RN*, which may also contribute to fine-tuning of IL1 signaling and risk conferred by variants at this locus. Together these data suggest that M2 macrophage-derived IL1A/B could be involved in endometriosis risk.

### Expression quantitative trait locus (eQTL) analysis links endometriosis risk SNPs to the expression of IL1A, IL1B and other candidate genes

Endometriosis risk variants in the chromosome 2q14.1 locus were mapped to eQTLs to identify evidence that genetically regulated expression differences in *IL1A* and/or *IL1B* are conferred by the same variants associated with endometriosis risk. Using the Fuma software and One1K single cell eQTL data we interrogated 139 eQTL datasets representing 37 different tissue types (Supplemental Table 3). Decreased expression of *IL1A* was associated with increased risk in 10 different tissue types (*P <* 10^-5^). Elevated endometriosis risk was associated with lower expression of *IL1B* in two myeloid cell types (monocytes, dendritic cells) and elevated expression in three data sets (Fig. 2e). The greatest number of associations were detected for *IL1A* (10 eQTLs). Five eQTL associations were detected for *IL1B* and *PSD4*. The number and strength of associations for *IL1A*, and the myeloid-specific associations for *IL1B* are further evidence to link these genes to endometriosis risk. To test whether the same variants are associated with risk and IL1A and/or IL1B expression, colocalization analyses were performed on a selection of tissue types. This revealed evidence of a shared causal variant between the endometriosis risk variants and a IL1A eQTL (Supplemental Table 4). The signals for IL1B look to be independent of the endometriosis risk association and are likely a consequence of linkage disequilibrium.

### M2 macrophages are central to paracrine signaling in endometriosis

CellChat ^30^ was used to identify potential signaling between cell types in endometriosis. In total 192 ligand-receptor pairs and 4,056 predicted connections were identified between cell types (P < 0.05, Supplemental Table 5), since different cell types can communicate through the same ligand-receptor pairs. Non-immune cells are predicted to form more outgoing than incoming links, while the opposite happens for immune cells (Fig. 3A). For example, endometrial-type stromal cells exhibit 439 outgoing and 99 incoming connections, while early exhausted T cells are involved in 124 outgoing and 385 incoming signals.

**Fig 3:**
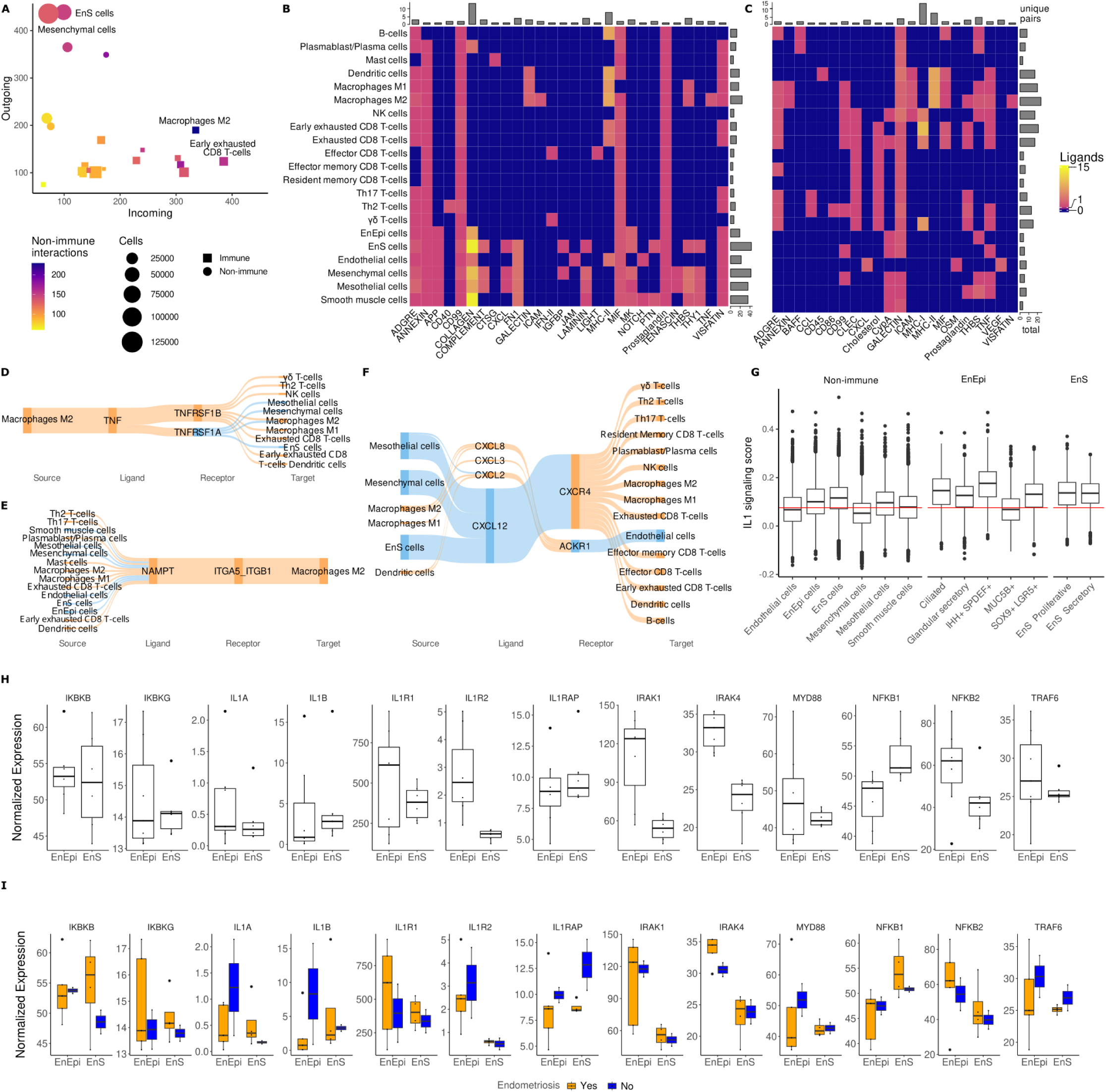
M2 macrophages play a central role in cell-cell communications in endometriosis. A) Predicted interactions for every cell cluster. B-C) pathways affected by B) incoming and, C) outgoing signals for M2 macrophages. D-F) Ligand-receptor pairs identified for D) TNF, E) Visfatin, F) CXCL. G) IL1 signaling expression in non-immune cell clusters and subclusters. H) Expression of interleukin-1 pathway genes in EnEpi and EnS. I) Expression of interleukin-1 pathway genes in EnEpi and EnS of women with and without a confirmed diagnosis of endometriosis. EnEpi, endometrial-type epithelium; EnS, endometrial-type stroma; NK, natural killer cells.

Non-immune cells tend to be more connected than immune cells, with M2 macrophages and early exhausted T cells particularly connected among immune cells, with 549 and 528 connections, respectively. Compared to other immune cells, M2 macrophages are exceptionally connected with non-immune cells, which form 229 interactions with M2 macrophages, contrasting to 157 connections for early exhausted T cells, the second most connected subset.

Interactions between non-immune cells and M2 macrophages are mostly incoming and concentrate on the collagen pathway (Fig. 3B-C). There are pathways affected by the same small set of LR pairs expressed indiscriminately across various cell types, like incoming CD99 and outgoing galectin, and pathways linked with distinct LR pairs affected in a small set of cell types like incoming collagen, incoming MHC-I and both incoming and outgoing MHC-II. Non-immune cells are under-represented among the recipients of outgoing interactions from M2 macrophages, with myeloid cells, exhausted T cells and NK cells being the predominant targets. All this suggests that M2 macrophages play a central role in the cross-talk between diagnostic and structural cells. Indeed M2 macrophages represent a communication bottleneck for a number of pathways including outgoing TNF (Fig. 3D) and incoming visfatin ^31^ signaling (Fig. 3E). In other instances, such as the CXCL pathway, myeloid cells, including M2 macrophages, participate in incoming and outcoming signaling with a variety of cell types, with specificity determined by the receptor and ligand molecules expressed (Fig. 3F)IL1 signaling was not identified in the CellChat analyses, likely due to low expression of the complex*IL1R1-IL1RAP*, which is the receptor form included in the database. We therefore performed a targeted survey of IL1 signaling among the non-immune cell types, searching for possible effector cells for the IL1A/B produced by M2 macrophages. Except MUC5+ endometrial-type epithelial cells, all endometrial-type epithelial (EnEpi) and stromal (EnS) subtypes surpass the mean expression of an IL1 signature among non-immune cell types, indicating that they are likely recipients of IL1 signaling. Endometrial-type stromal cells and the IHH+SPDEF+ subset of endometrial-type epithelial cells exhibit the highest average expression of the IL1 signature among non-immune cell types (Fig. 3G, Supplementary Fig. 2). To evaluate the expression of IL1 signaling genes in cells from the eutopic endometrium from cases and controls, bulk-RNA sequencing data were generated from freshly isolated primary endometrial epithelia (EnEpi) and endometrial stroma (EnS). Bulk RNA-sequencing provides more depth and can identify more lowly expressed genes compared to single-cell sequencing.

Analysis of marker genes for IL1 signaling revealed positive expression of the 13 genes examined (Figure 3H). The pattern of expression for candidate genes between epithelial and stromal cells appeared similar with more variation within EnEpi. IL1R2 and signaling intermediaries *IRAK1* and *IRAK4* show the highest significant difference between EnEpi and EnS. *IL1R1* has high expression in both cell types, but the associated protein *IL1RAP* is relatively low, consistent with the single cell data. While *IL1A* and *IL1B* were positive in both cell types their expression was low compared to *IL1R1*. (Fig. 3I). There was also a tendency for eutopic endometrium from endometriosis cases to express higher *IL1R1* and lower *IL1A* and *IL1B*.

### IL1A/B treatment of endometrial organoids *in vitro* disrupts epithelial differentiation

When maintained in Matrigel with defined growth factors (see Methods), EnEpi cells co-cultured with EnS or endometrial mesenchymal stem cells spontaneously create complex assembloids that closely mimic human endometrial tissues ^32^. Epithelial cells were isolated from menstrual fluid from six patients with suspected endometriosis, five had endometriosis confirmed through laparoscopic surgery, and the sixth did not undergo surgery, leaving endometriosis status unconfirmed. Stromal cells were isolated from endometrial biopsies from 8 different patients (4 with endometriosis and 4 without endometriosis) and the two cell types were co-cultured to establish assembloids. After single-cell RNA-sequencing of the assembloids, cell identity was transferred from the endometriosis cell atlas, stratifying epithelial cells into ciliated and secretory subsets (Fig. 4A and Supplementary Fig. 3). IL1 signaling was elevated in ciliated cells relative to non-ciliated epithelia, although IL1A and IL1B are under-expressed (Fig. 4B,C). IL1 signaling was positively correlated with estrogen signaling in ciliated cells only (Fig. 4D, P-value = 0.017), consistent with the known role of estrogen on ciliogenesis ^33^ and the substantial overlap in the transcriptional targets of estrogen and IL1 (89.47% of the IL1 signaling genes are also estrogen regulated ^34^).

**Fig. 4:**
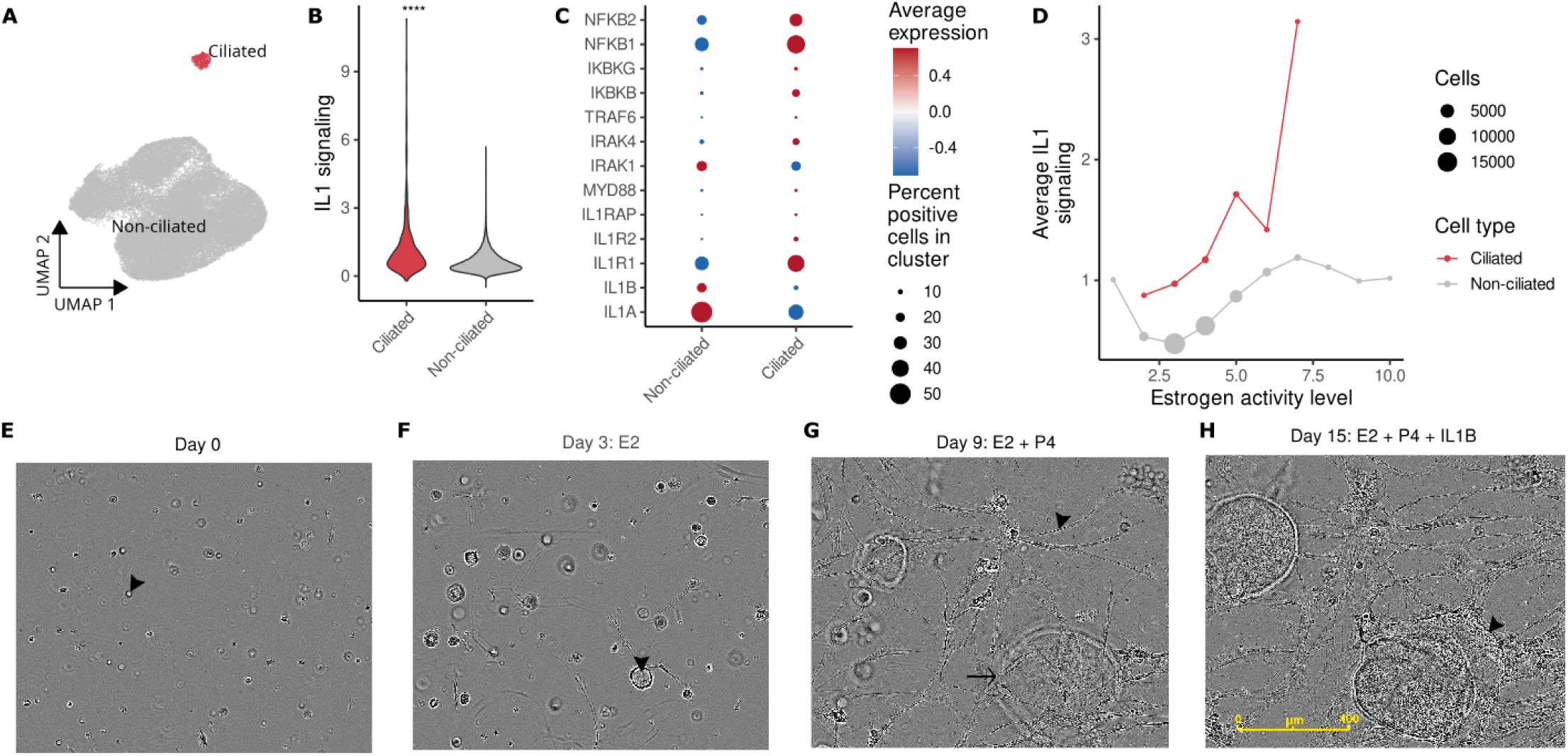
Stimulation of interleukin-1 signaling disrupts the organization of human endometrial organoids. A) UMAP representation of single cell transcriptomes for 39,568 endometrial epithelial cells from organoids. B) IL1 signaling for ciliated and non-ciliated cells. T-test P - value < 0.0001(****). C) Expression of selected markers from IL1 signaling pathway. D) Correlation between estrogen activity and IL1 signaling. E) Single cell seeding of epithelial and mesenchymal stem cells. Arrowhead denotes a representative single cell. F) Treatment with estradiol (E2) and cell proliferation. Arrowhead shows a representative proliferating organoid. G) Treatment with E2 in combination with progesterone (P4), stromal cell decidualization (arrowhead) and epithelial organoid expansion (arrow). H) Treatment with IL1B results in disordered epithelial differentiation (arrowhead).

To test the impact of IL1B exposure *in vitro*, single cell suspensions of endometrial epithelial cells and endometrial mesenchymal stem cells were seeded into Matrigel and cultured under conditions to facilitate the generation of 3-dimensional *in vitro* endometrial tissue (Fig. 4E). To mimic endometrial development and changes across the menstrual cycle, assembloids were treated with estradiol (E2), which stimulated cell proliferation, characteristic of proliferative (follicular) phase endometrium (Fig. 4F). A combination of E2 with progesterone (P4) resulted in a transition to elongated decidualised mesenchymal stromal cell knots and expansion of epithelial glandular organoid structures (Fig. 4G), that mimic the morphology and structure of mature secretory stage endometrium. Treatment of mature, decidualized assembloids with IL1B resulted in a drastic alteration in cellular morphology, with extensive reorganization of the epithelial glandular structures. Disrupted glandular differentiation resulted in epithelial spread and infiltration into the surrounding decidualized mesenchyme (Fig. 4H, Supp. video 1)

### IL1R antagonist therapy reduces pain sensitivity *in vivo*

Since the scDRS and *in vitro* results support a central role for IL1 signaling in endometriosis pathogenesis, we evaluated the effect of using anakinra to disrupt intercellular IL1 signaling in a validated mouse model of endometriosis-associated pain ^35^. Anakinra is a pharmacologically optimized recombinant version of interleukin-1 receptor antagonist, the product of the *IL1RN* gene. Endometriosis-like lesions were induced in mice by injecting dissociated uterine fragments from estrogen-primed donor animals into the peritoneal cavity of syngeneic recipients. A blinded investigator measured evoked pain using von Frey fibers on day 0 and weekly thereafter. On day 29 animals were block randomized to treatment groups and treated for 4 weeks. At the end of the experiment a blinded investigator measured evoked pain, spontaneous pain (abdominal squashing and abdominal contortions), and lesion size and number. Anakinra treatment results in a significant, dose-dependent decrease in spontaneous pain and evoked pain (Fig. 5A-C).

**Fig. 5:**
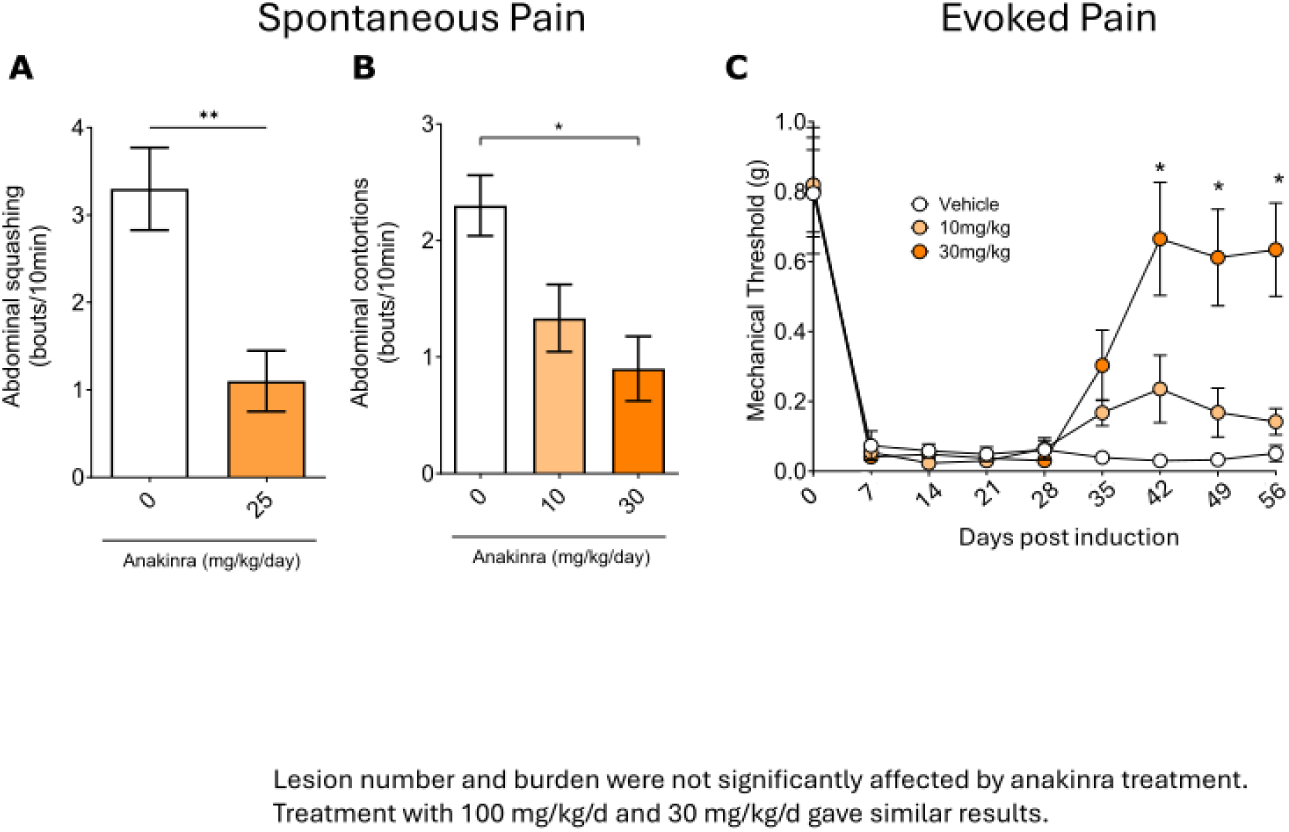
The effect of an IL-1 inhibitor on endometriosis-associated pain in a mouse model. Endometriosis-like lesions were induced in C57BL/6J mice, allowed to grow for 4 weeks, and treated with anakinra at the indicated dose. Spontaneous pain was decreased following 4 weeks of treatment. A) Abdominal squashing B) Abdominal contortions. C) Evoked pain measured using von Frey fibers on the abdomen, was also decreased in a dose-dependent manner. *P-value < 0.05 by analysis of variance (ANOVA). Investigators were blinded to group allocation until all analysis was complete.

### M2 macrophages explain genetic interactions with immune and pain conditions

An extensive body of literature documents epidemiologic and genetic links between endometriosis and inflammatory conditions. Asthma (odds ratio [OR] = 1.35, 95% confidence interval [CI] = 0.97-1.88), chronic fatigue syndrome and/or fibromyalgia (OR = 5.81, 95% CI = 1.89-17.9), mononucleosis (OR = 1.75, 95% CI = 1.14-2.68), allergy (OR = 1.76, 95% CI = 1.32-2.36) and rheumatoid arthritis (IRR = 1.31, 95% CI: 1.05-1.64) were associated with endometriosis ^10,36–40^. We have recently also identified a significant genetic correlation and causal relationship between genetic liability to inflammatory gastrointestinal disorders and endometriosis risk, and evidence for a bidirectional causal relationship between endometriosis and irritable bowel syndrome^41^. To check for other risk associations of the endometriosis microenvironment, summary statistics from GWAS for 16 traits (Supplemental table 6) were used to calculate the scDRS (Supplemental table 7). B cells (but not plasma cells), dendritic cells, M1 and M2 macrophages as well as functional T cell subsets, were all associated with arthritis, asthma and irritable bowel syndrome (Fig. 6). Dendritic cells, M1 and M2 macrophages and endometrial-type epithelium were all associated with abdominal and pelvic pain but not back pain.

**Fig. 6:**
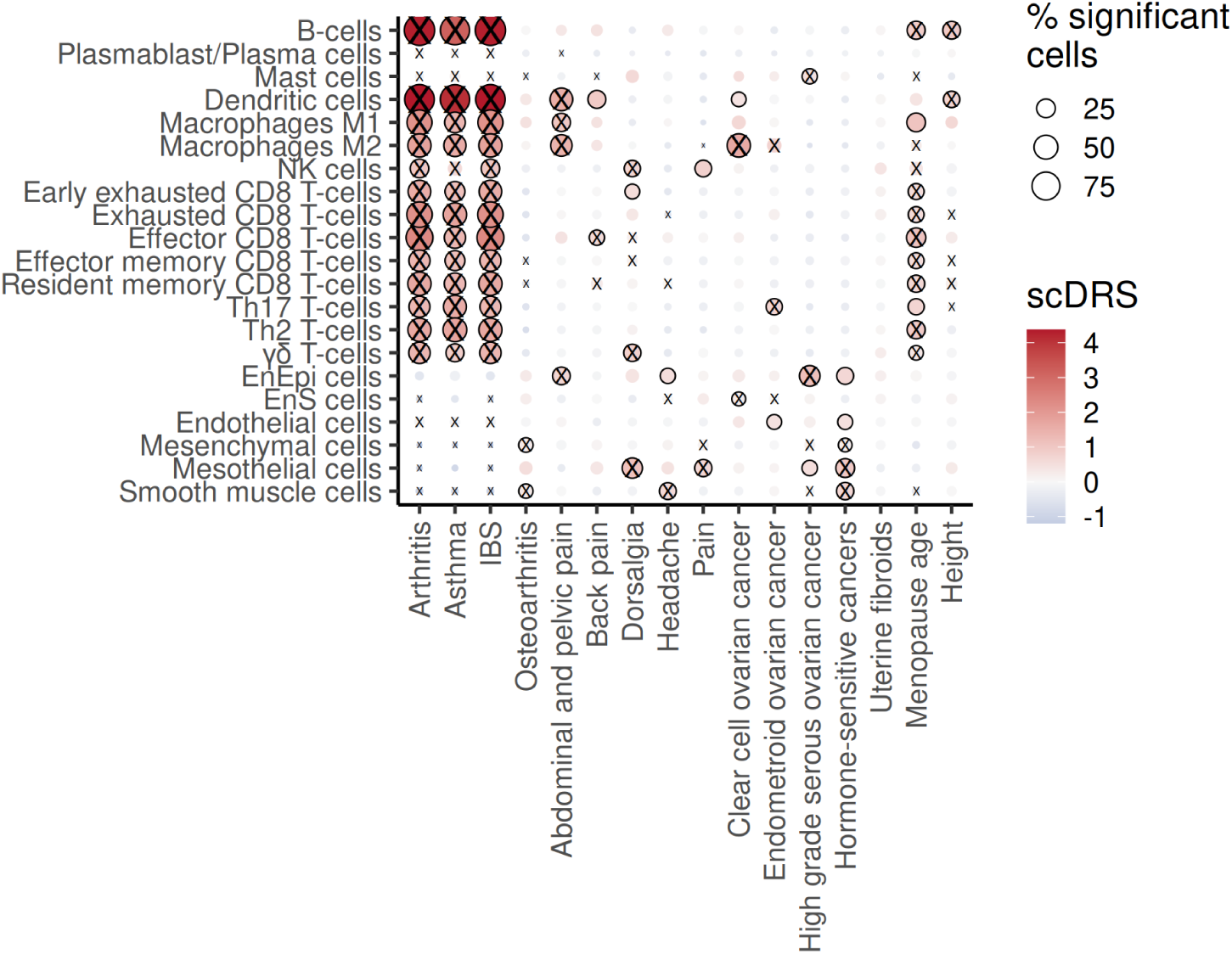
scDRS identifies cell types that may explain associations with traits that have shared germline genetics with endometriosis. scDRS scores for other traits that share risk genetics with endometriosis. A cross symbol indicates significant heterogeneity detected.

Endometriosis is associated with risk of clear cell and endometrioid ovarian cancer, but links with high-grade serous subtype are less consistent ^4,42^. M2 macrophages and endometrial-type stroma were associated with clear cell ovarian cancer and are heterogenous in endometrioid ovarian cancer (Fig. 6). Endometrial-type epithelium, mesothelial cells and other structural cells had associations with high-grade serous ovarian cancers and/or hormone-sensitive cancers (postmenopausal breast, uterine, ovary, prostate, and thyroid), likely reflecting a shared cell-intrinsic component to development of these cancers. Somewhat unexpectedly, cell types associated with age at menopause were similar to those associated with arthritis, asthma and irritable bowel syndrome, suggesting immune cells play a greater role in the inherited component of age at menopause than previously thought.

## Discussion

Endometriosis has myriad long-term negative effects on women’s health. There are no known highly penetrant germline mutations that contribute to endometriosis risk; but a substantial component of endometriosis risk is inherited, with heritability estimated at 47-51% from twin studies ^43,44^. Most of the inherited risk for endometriosis is due to the combinatorial effects of common mild-risk variants. So far, 42 risk loci have been identified for endometriosis^10,11,13,45,46^. Interpretation of post-GWAS studies to date have focused on the role of endometrial-type epithelial cells, however given the central role of the microenvironment in disease pathogenesis, this likely overlooks a critical aspect of risk. Here, by integrating single cell gene expression profiles of endometriosis with GWAS data for endometriosis, M2 macrophages, dendritic cells and endothelial cells were associated with endometriosis risk, suggesting these cells play a role in disease susceptibility and development.

Evidence for genetic variants proximal to the interleukin 1A gene locus were among the earliest risk regions identified for endometriosis ^27,28^. Of all the cytokine families, IL1 is most associated with damaging inflammation ^47–49^ and has been predicted to modulate endometriosis symptomatology ^50^. In this study, IL1A/IL1B signaling was identified as a major effector driving the association between M2 macrophages and endometriosis.

Somewhat unexpectedly, in eQTL analyses, lower IL1A expression was associated with increased risk, suggesting that muted inflammatory responses may be protective against developing endometriosis. However the eQTL results should be interpreted with caution as IL1A and IL1B are known to have highly context-specific specific effects and their roles in endometriosis development and symptomatology are not yet completely defined ^51^. Computational inference of cell-cell communications define M2 macrophages as a major hub of signaling between immune and non-immune cells, with IL1 signatures activated in endometrial-type stroma and epithelial subsets, identifying these diagnostic cells within lesions as effectors of IL1 signaling in endometriosis. IL1A and IL1B both signal through the interleukin receptor (*IL1R1*) to regulate NFΚB, cJUN and p38 MAPK signaling, inducing the expression of a host of genes including IL8 ^52^. Expression of IL8 (encoded by the *CXCL8* gene) was also associated with endometriosis single cell disease risk scores, suggesting multiple risk variants may converge to modulate different components of the IL1 pathway. A positive correlation with NF-κB and TNF-α signaling and risk score supports previous observations of a triggered inflammatory response in endometriosis macrophages ^53^.

*In vivo* targeting of IL1 signaling reduced pain symptoms in mice with induced endometriosis, consistent with a recent pilot trial in humans which reported reduced dysmenorrhea (pelvic pain with menstruation) and improved quality of life in patients receiving 3 months of Anakinra (IL1 receptor antagonist) treatment compared to placebo ^54^. Moreover translational studies link IL1B protein expression to neurogenesis as well as stromal cell migration and invasion ^48,55^, suggesting IL1 receptor antagonist treatment could have pleiotropic effects on lesions and rendering this a particularly attractive non-hormonal therapeutic strategy.

M2 macrophages and dendritic cells were also associated with immune-related traits that have genetic associations with endometriosis ^10,56^, indicating that pleiotropic multi-cellular mechanisms may explain genetic overlap between pain and inflammatory conditions. Expression QTL analyses identified associations with *IL1A* and *IL1B* in the blood, suggesting a model whereby macrophages, differentiated from monocytes that circulate in the blood, contribute to risk of multiple inflammatory traits including endometriosis ^20^. In this pleiotropic model, receptor expression occurs in a disease-relevant cell type - epithelial cells in endometriosis (we note that lung alveolar epithelial cells also express *IL1R1* and variants proximal to *IL1R1* are associated with risk of asthma).

The main limitation of this study is the dependency on gene expression; epigenome and functional studies will be needed to causally link endometriosis-associated genetic variation to M2 macrophage function and modified IL1 signaling. Sample size of the gene expression dataset could also bias observations. However, convergence of evidence from orthogonal approaches on IL1A/B, support continued investigation of IL1R therapy for endometriosis ^54^, where new non-hormonal therapeutic options are urgently needed.

## Methods

### Single cell endometriosis profiles, immune cell annotation

Data from GSE213216 was used for analysis ^8^. 101,217 T/natural killer T-cells, 27,436 myeloid cells, 8,278 B/plasma cells and 1,687 mast cells were included in the analysis, plus 10,179 additional cells that originally clustered with epithelial and mesenchymal cells but were removed from analysis due to the absence of epithelial or mesenchymal cell markers and positive expression of canonical immune markers. Four samples annotated as ‘endometriosis-free’ in our initial report ^8^ were reclassified as peritoneal endometriosis due to the detection of endometriosis-type epithelium and/or stroma in those specimens.

CD45 (*PTPRC*) plus additional immune markers were used to move cluster 23, 74, and 101 (cluster number from Fonseca *et al,* ^8^ these clusters were originally subgrouped with mesenchymal cells but excluded during reclustering) to the T/NK cell group. Any other clusters that express immune markers but low expression of *PTPRC* (ie, certain myeloid) were also selected to be moved (cluster 103 to myeloid group). From the cells removed from the epithelial group, cluster 53 were moved to the myeloid cell group.

All immune cell clusters (B/plasma cells, mast cells, myeloid cells, T/NK cell clusters) were reclustered at resolution 0.6, resulting in 24 clusters. Immune clusters were annotated using canonical markers for T cells (*CD3D*, *CD8A*, *CD4*; 87,953 cells), natural killer cells (*KIR2DL4*, *NCAM1*; 5,977 cells), myeloid cells (*CD14*, *LYZ*, *CD68*; 15,483 cells), B cells, plasmablast/plasma cells (*MS4A1*, *CD79A*, *JCHAIN*, *SDC1*, *CD27*, 5,274 and 1,771 cells) and mast cells (*KIT*, *TPSB2*; 1,645 cells).

Seurat clusters expressing markers for the same cell type were merged, i.e. 5 clusters with markers for Effector Memory CD8 T-cells are all labeled the same, resulting in 15 distinct functional populations.

Unless otherwise specified, analysis were performed using R programming language 4.3.

### scDRS analysis

GWAS summary statistics were downloaded for endometriosis and 16 other traits ^57–65^. Trait sample size and ID are listed in Supplementary Table 6. Sample size was maintained over 100,000 to follow software recommendations, except for arthritis, which only includes 97,173 subjects. When missing, reference SNP ids were obtained using intersect bedtools between GWAS coordinates and all GRCh37 SNPs. Reference SNP IDs are needed by MAGMA ^66^ to get a score of disease association per gene. The scDRS software takes as input MAGMA scores for the top 1,000 associated genes and outputs a score and P-value per cell. Afterwards, group analysis was tested with the downstream command of the scDRS software for cell type, disease class and stage. rASRM stages 1 and 2 are relabeled as low stage, while 3 and 4 become high. All scDRS steps were run with default parameters, without including covariates.

Spearman correlation was estimated between the expression pattern of every gene and used for ranking and GSEA functional enrichment against the database of reactome pathways.

In order to complement scDRS heterogeneity analysis, pathway activity was estimated using the progeny R package ^23^. Activity estimates were binned and contrasted with the scDRS, per cell type.

### eQTL Mapping

FUMA v1.6.1 ^67^ was used to map cis-eQTLs to endometriosis GWAS variants in the chromosome 2q14.1 locus using a GWAS P-value threshold of p<5x10^-8^ to select SNPs and FDR<0.05 to select SNP-Gene associations. The locus was defined as ±1MB from the lead GWAS SNPs (rs3783513 & rs10167914). Similarly, to investigate if GWAS variants in the region have been associated with regulation of expression in immune cell types we mapped significant (p<5x10-8) GWAS variants in the chromosome 2q14.1 locus to cis-eQTLs identified using scRNA-seq in the ONEK1K study ^68^ consisting of 26,597 independent eQTLs (p<0.05) across 14 immune cell types ^68^. The most significant eQTL for each gene in each tissue/cell type was plotted.

### Analysis of bulk RNA sequencing data from isolated epithelial and stromal cells

Using freshly isolated endometrial epithelial and stromal cells from women with and without endometriosis and performing bulk RNA-sequencing analysis we assessed the expression of IL1 signaling markers, which included *IL1A, IL1B, IL1R1, IL1R2, IL1RAP, MYD88, IRAK1, IRAK4, TRAF6, IKBKB, IKBKG, NFKB1, NFKB2*.

Markers are taken from reactome pathway R-HSA-9020702. Differential gene expression analysis was performed after TMM normalization, with a linear regression model and included cell type, cycle stage at isolation and presence of endometriosis as covariates in the model.

### *In vitro* modeling of endometrium and endometriosis

Complex *In vitro* models of endometrium were established to assess IL1/IL1R expression in mature, refluxed endometrium by generating organoids from menstrual fluid and combining with endometrial stromal cells to introduce impacts derived from cellular interaction. The impact of hormonal exposure on endometrial maturation was also assessed *in vitro* by initiating assembloid models with endometrial stem cells and exogenously mimicking hormonal stimulated endometrial development through the proliferative, secretory and subsequent “ectopic” stage.

Human menstrual fluid was collected from premenopausal women using a silicone menstrual cup (Lunette, Juupajoki, Finland) as described previously ^69^. Informed consent was obtained for each patient prior to surgery. Epithelial cell adhesion molecule (EPCAM) magnetic beads (CELLection Epithelial Enrichment kit, Invitrogen) were used to enrich epithelial cells that were subsequently seeded into Matrigel (7,000 clusters per well) and maintained in organoid-specific media ^70^. To generate allogenic assembloid co-cultures endometrial stromal cells were obtained from endometrial biopsies from 8 women (4 with and 4 without endometriosis), separated and cultured as described previously ^71^ and subsequently mixed at a ratio of 1:1 of organoid to stromal cell, with a subsequent ratio of 1:7 of cell mixture with Matrigel to establish mature endometrial *in vitro* models.

To generate hormonally stimulated endometrial assembloids, endometrial biopsies were dissociated to isolate single epithelial progenitor cells and mesenchymal stem cells. Endometrial mesenchymal stem cells were collected from the mesenchymal compartment and induced into a stem cell state through seeding at a sparse density (5,000 cells/cm^2^) to achieve clonal growth from individual cells and maintained (> 1 month) in specialized stem cell media (DMEM/F12, 1µm A83-01, 10ng/ml EGF and 10ng/ml FGF2). Mesenchymal stem cell capacity was confirmed through the ability to differentiate cells into alternate mesodermal lineages of osteoblasts, chondrocytes and adipose cells. Single epithelial progenitors and mesenchymal stem cells were combined in a 1:1 ratio, seeded into Matrigel and maintained in organoid media, allowing co-ordinate maturation of both epithelial and mesenchymal cells ^72,73^. To simulate hormone exposure and menstrual stage cycles immature assembloids were exposed to 10nM estrogen (day 0-5; proliferative stage), with a subsequent 10nM Estrogen, 1um Progesterone and 1 um cAMP (day 5-12; secretory stage). To mimic exposure to an inflammatory ectopic environment hormonally-matured assembloids were exposed to 10ng/µL IL1B (day 12-19). Cell growth and morphology was monitored via growth in the Incucyte Imaging system (Sartorius).

#### Annotation of organoid cells

The default Seurat pipeline for dataset integration was run to transfer labels from the single cell patient data to the organoids dataset, including SCT normalizing and anchor finding based on variable genes. Two levels of cell type identification were attained by transferring cell type labels and endometrial epithelial cell subtype labels. The 2.81% of cells with prediction score below 0.75 were discarded as non-identifiable. Quality of the results was assessed based on expression of published cell type markers.

### In vivo assays

#### Induction of Endometriosis-like lesions

Lesions were induced as described previously ^35^. Briefly, after at least one week of acclimatization, donor mice received a subcutaneous injection of 3 μg/mouse estradiol benzoate to stimulate the growth of the endometrium. Four days later, the uteri of the donor mice were dissected into a Petri dish containing Hank’s Balanced Salt Solution (HBSS, Thermo Fisher Scientific, Waltham, MA, USA) and split longitudinally with a pair of scissors. Uterine horns from each donor mouse were minced with scissors and scalpel one at the time, ensuring that the maximal diameter of each fragment was consistently smaller than 1 millimeter (mm). Each dissociated uterine horn was then injected intraperitoneally using an 18G needle (cat #305185 Thin wall, BD, Franklin Lakes, NJ, USA) into a recipient mouse in 500 µL of HBSS. One donor mouse was used for every two endometriosis mice.

#### Study design

Block randomization was used to randomize subjects into groups of 10 mice each. Experiments were powered for determining responses to pain but not lesion size. Mice were treated by injection with anakinra (Boston Children’s Hospital Pharmacy) in sterile saline (0, 25mg/kg/day; 0, 10, 30, 100mg/kg/day) starting at day 29 and ending on day 56. The investigators were blinded to the treatment groups in all testings until the end of the experiment and analysis.

#### Behavioral testing

Mice were allowed to habituate to the apparatus for at least 2h and during three consecutive days before the beginning of measurements. After habituation, baseline measurements were obtained one day prior to the induction of endometriosis. Pain intensity to a mechanical stimulus (mechanical hyperalgesia) in the abdominal region was measured using von Frey filaments. The experimenter was trained, and care was taken not to stimulate the same point consecutively, and the stimulation of the external genitalia was avoided. A jump or paw flinch was considered a withdrawal response ^74^. The mechanical threshold was determined by the up and down method starting with 0.4g filament and calculated using the open-source software Up-Down Reader ^75^.

For spontaneous abdominal pain measurements, stretching the abdomen (abdominal contortions), and squashing of the lower abdomen against the floor were quantified as previously described ^74,76,77^. For abdominal contortions, mice were placed in individual chambers in a temperature-controlled (29°C) glass plate and the number of abdominal contortions was quantified with a positive response consisting of a contraction of the abdominal muscle together with stretching of hind limbs. For abdominal squashing, the number of times the mice pressed the lower abdominal region against the floor was determined. In all testing, the investigators were blinded to the treatments.

#### Statistical analysis

Results are presented as mean ± SEM. Data were analyzed using the software GraphPad Prism version 10.1.2 (GraphPad Software, San Diego, CA, USA). Two-way repeated measure analysis of variance (ANOVA), followed by Tukey’s *post hoc*, was used to analyze data from mechanical hyperalgesia. An unpaired T-test was used to analyze data from experiments with a single time point. For the percentage of mice with visible lesions, statistical analysis was estimated by the Kaplan-Meier method followed by the logrank test. For all analyses, statistical differences were considered significant when p < 0.05.

#### Ethics approval

All mouse work was performed according to protocols approved by the Institutional Animal Care and Use Committee (IACUC) at Boston Children’s Hospital (protocols 19-12-4054R and 00001816). The work followed the Guide for the Care and Use of Laboratory Animals and all of the regulatory protocols set forth by the Boston Children’s Hospital Animal Resources at Children’s Hospital (ARCH) facility.

## Supplemental information

Document S1. Figures S1–S3

Document S2. Excel file containing supplemental tables

Document S3. Seeding of epithelial and mesenchymal stem cells, related to Figure 4E.

Document S4. Treatment of assembloids with estradiol (E2) and cell proliferation, related to Figure 4F.

Document S5. Treatment of assembloids with E2 in combination with progesterone (P4), related to figure 4G

Document S6. Treatment of assembloids with IL1B, related to figure 4H

Document S7. Table S7 containing the scDRS results

Video S1. Assembloid growth and response to IL1B treatment, related to Figure 4

## Data availability

The single cell endometriosis atlas data is publicly available at GSE213216. The rest of the datasets are in the process of submission.

## Code availability

The code used for figure generation, scDRS and CellChat analysis is provided in: https://github.com/lawrenson-lab/scDRS_endometriosis

## Acknowledgements

Specimens were collected as part of the Biologic and Epidemiologic Markers of Endometriosis (BEME) study and Women’s Cancer Biobank within the Department of Obstetrics and Gynecology at Cedars-Sinai Medical Center. We thank Dr Caitlin Filby for generating the menstrual fluid organoids. We wholeheartedly thank all the patients who donated the specimens used in this study.

## Author contributions

Conceptualization (SO, KL, GWM, MSR, BM); Formal analysis (SO); Investigation (SO, FSR-O, SM, SC, PY); Data curation (MH); Resources (SS, CEG); Writing - original draft (SO, KL); Writing - review and editing (all authors); Supervision (KL); Funding acquisition (KL, MSA, GWM, CEG, MSR).

## References

1. C. L. Pearce, C. Templeman, M. A. Rossing, A. Lee, A. M. Near, P. M. Webb, C. M. Nagle, J. A. Doherty, K. L. Cushing-Haugen, K. G. Wicklund, J. Chang-Claude, R. Hein, G. Lurie, L. R. Wilkens, M. E. Carney, M. T. Goodman, K. Moysich, S. K. Kjaer, E. Hogdall, A. Jensen, Ovarian Cancer Association Consortium, Lancet Oncol. 2012, 13, 385–394.

2. M. Kvaskoff, Y. Mahamat-Saleh, L. V. Farland, N. Shigesi, K. L. Terry, H. R. Harris, H. Roman, C. M. Becker, S. As-Sanie, K. T. Zondervan, A. W. Horne, S. A. Missmer, Hum. Reprod. Update 2021, 27, 393–420.

3. T. Zhang, C. De Carolis, G. C. W. Man, C. C. Wang, Autoimmun. Rev. 2018, 17, 945–955.

4. J. Vallvé-Juanico, S. Houshdaran, L. C. Giudice, Hum. Reprod. Update 2019, 25, 564–591

5. M. A. S. Fonseca, M. Haro, K. N. Wright, X. Lin, F. Abbasi, J. Sun, L. Hernandez, N. L. Orr, J. Hong, Y. Choi-Kuaea, H. M. Maluf, B. L. Balzer, A. Fishburn, R. Hickey, I. Cass, H. S. Goodridge, M. Truong, Y. Wang, M. D. Pisarska, H. Q. Dinh, K. Lawrenson, Nat. Genet. 2023, 55, 255–267.

6. Y. Tan, W. F. Flynn, S. Sivajothi, D. Luo, S. B. Bozal, M. Davé, A. A. Luciano, P. Robson, D. E. Luciano, E. T. Courtois, Nat. Cell Biol. 2022, 24, 1306–1318.

7. M. J. Zhang, K. Hou, K. K. Dey, S. Sakaue, K. A. Jagadeesh, K. Weinand, A. Taychameekiatchai, P. Rao, A. O. Pisco, J. Zou, B. Wang, M. Gandal, S. Raychaudhuri, B. Pasaniuc, A. L. Price, Nat. Genet. 2022, 54, 1572–1580.

8. E. M. Weeks, J. C. Ulirsch, N. Y. Cheng, B. L. Trippe, R. S. Fine, J. Miao, T. A. Patwardhan, M. Kanai, J. Nasser, C. P. Fulco, K. C. Tashman, F. Aguet, T. Li, J. Ordovas-Montanes, C. S. Smillie, M. Biton, A. K. Shalek, A. N. Ananthakrishnan, R. J. Xavier, A. Regev, H. K. Finucane, Nat. Genet. 2023, 55, 1267–1276.

9. K. A. Jagadeesh, K. K. Dey, D. T. Montoro, R. Mohan, S. Gazal, J. M. Engreitz, R. J. Xavier, A. L. Price, A. Regev, Nat. Genet. 2022, 54, 1479–1492.

10. N. Rahmioglu, S. Mortlock, M. Ghiasi, P. L. Møller, L. Stefansdottir, G. Galarneau, C. Turman, R. Danning, M. H. Law, Y. Sapkota, P. Christofidou, S. Skarp, A. Giri, K. Banasik, M. Krassowski, M. Lepamets, B. Marciniak, M. Nõukas, D. Perro, E. Sliz, K. T. Zondervan, Nat. Genet. 2023, 55, 423–436.

11. J. N. Painter, C. A. Anderson, D. R. Nyholt, S. Macgregor, J. Lin, S. H. Lee, A. Lambert, Z. Z. Zhao, F. Roseman, Q. Guo, S. D. Gordon, L. Wallace, A. K. Henders, P. M. Visscher, P. Kraft, N. G. Martin, A. P. Morris, S. A. Treloar, S. H. Kennedy, S. A. Missmer, K. T. Zondervan, Nat. Genet. 2011, 43, 51–54.

12. N. Rahmioglu, D. R. Nyholt, A. P. Morris, S. A. Missmer, G. W. Montgomery, K. T. Zondervan, Hum. Reprod. Update 2014, 20, 702–716.

13. Y. Sapkota, V. Steinthorsdottir, A. P. Morris, A. Fassbender, N. Rahmioglu, I. De Vivo, J. E. Buring, F. Zhang, T. L. Edwards, S. Jones, D. O, D. Peterse, K. M. Rexrode, P. M. Ridker, A. J. Schork, S. MacGregor, N. G. Martin, C. M. Becker, S. Adachi, K. Yoshihara, D. R. Nyholt, Nat. Commun. 2017, 8, 15539.

14. M. L. Freedman, A. N. A. Monteiro, S. A. Gayther, G. A. Coetzee, A. Risch, C. Plass, G. Casey, M. De Biasi, C. Carlson, D. Duggan, M. James, P. Liu, J. W. Tichelaar, H. G. Vikis, M. You, I. G. Mills, Nat. Genet. 2011, 43, 513–518.

15. S. Mortlock, S. Houshdaran, I. Kosti, N. Rahmioglu, C. Nezhat, A. F. Vitonis, S. V. Andrews, P. Grosjean, M. Paranjpe, A. W. Horne, A. Jacoby, J. Lager, J. Opoku-Anane, K. C. Vo, E. Manvelyan, S. Sen, Z. Ghukasyan, F. Collins, X. Santamaria, P. Saunders, L. Giudice, Commun. Biol. 2023, 6, 780.

16. S. Mortlock, R. I. Kendarsari, J. N. Fung, G. Gibson, F. Yang, R. Restuadi, J. E. Girling, S. J. Holdsworth-Carson, W. T. Teh, S. W. Lukowski, M. Healey, T. Qi, P. A. W. Rogers, J. Yang, B. McKinnon, G. W. Montgomery, Hum. Reprod. 2020, 35, 377–393.

17. S. Mortlock, R. Restuadi, R. Levien, J. E. Girling, S. J. Holdsworth-Carson, M. Healey, Z. Zhu, T. Qi, Y. Wu, S. W. Lukowski, P. A. W. Rogers, J. Yang, A. F. McRae, J. N. Fung, G. W. Montgomery, Clin. Epigenetics 2019, 11, 49.

18. J. N. Fung, J. E. Girling, S. W. Lukowski, Y. Sapkota, L. Wallace, S. J. Holdsworth-Carson, A. K. Henders, M. Healey, P. A. W. Rogers, J. E. Powell, G. W. Montgomery, Hum. Reprod. 2017, 32, 893–904.

19. N. Rahmioglu, G. W. Montgomery, K. T. Zondervan, Womens Health (Lond Engl) 2015, 11, 577–586.

20. V. Fattori, T. H. Zaninelli, F. S. Rasquel-Oliveira, O. K. Heintz, A. Jain, L. Sun, M. L. Seshan, D. Peterse, A. E. Lindholm, R. M. Anchan, W. A. Veri, M. S. Rogers, BioRxiv 2024.

21. V. Steinthorsdottir, G. Thorleifsson, K. Aradottir, B. Feenstra, A. Sigurdsson, L. Stefansdottir, A. M. Kristinsdottir, F. Zink, G. H. Halldorsson, N. Munk Nielsen, F. Geller, M. Melbye, D. F. Gudbjartsson, R. T. Geirsson, U. Thorsteinsdottir, K. Stefansson, Nat. Commun. 2016, 7, 12350.

22. K. B. Zutautas, D. J. Sisnett, J. E. Miller, H. Lingegowda, T. Childs, O. Bougie, B. A. Lessey, C. Tayade, Front. Immunol. 2023, 14, 1089098.

23. M. Schubert, B. Klinger, M. Klünemann, A. Sieber, F. Uhlitz, S. Sauer, M. J. Garnett, N. Blüthgen, J. Saez-Rodriguez, Nat. Commun. 2018, 9, 20.

24. V. Lac, L. Verhoef, R. Aguirre-Hernandez, T. M. Nazeran, B. Tessier-Cloutier, T. Praetorius, N. L. Orr, H. Noga, A. Lum, J. Khattra, L. M. Prentice, D. Co, M. Köbel, V. Mijatovic, A. F. Lee, J. Pasternak, M. C. Bleeker, B. Krämer, S. Y. Brucker, F. Kommoss, M. S. Anglesio, Hum. Reprod. 2019, 34, 69–78.

25. T. Liu, L. Zhang, D. Joo, S.-C. Sun, Signal Transduct. Target. Ther. 2017, 2, 17023.

26. A. Badie, K. Saliminejad, I. Salahshourifar, H. R. Khorram Khorshid, Gynecol. Endocrinol. 2020, 36, 135–138.

27. Y. Sapkota, S.-K. Low, J. Attia, S. D. Gordon, A. K. Henders, E. G. Holliday, S. MacGregor, N. G. Martin, M. McEvoy, A. P. Morris, A. Takahashi, R. J. Scott, M. Kubo, K. T. Zondervan, G. W. Montgomery, D. R. Nyholt, Hum. Reprod. 2015, 30, 239–248.

28. S. Adachi, A. Tajima, J. Quan, K. Haino, K. Yoshihara, H. Masuzaki, H. Katabuchi, K. Ikuma, H. Suginami, N. Nishida, R. Kuwano, Y. Okazaki, Y. Kawamura, T. Sasaki, K. Tokunaga, I. Inoue, K. Tanaka, J. Hum. Genet. 2010, 55, 816–821.

29. S. Jin, C. F. Guerrero-Juarez, L. Zhang, I. Chang, R. Ramos, C.-H. Kuan, P. Myung, M. V. Plikus, Q. Nie, Nat. Commun. 2021, 12, 1088.

30. A. M. Krasnyi, A. A. Sadekova, T. Y. Smolnova, V. V. Chursin, N. A. Buralkina, V. D. Chuprynin, E. Yarotskaya, S. V. Pavlovich, G. T. Sukhikh, Int. J. Mol. Sci. 2022, 23, 10361.

31. T. M. Rawlings, K. Makwana, M. Tryfonos, E. S. Lucas, Reprod. Fertil. 2021, R85–R101.

32. S. Haider, M. Gamperl, T. R. Burkard, V. Kunihs, U. Kaindl, S. Junttila, C. Fiala, K. Schmidt, S. Mendjan, M. Knöfler, P. A. Latos, Endocrinology 2019, 160, 2282–2297.

33. O. Liska, B. Bohár, A. Hidas, T. Korcsmáros, B. Papp, D. Fazekas, E. Ari, Database (Oxford) 2022, 2022, baac083.

34. V. Fattori, N. S. Franklin, R. Gonzalez-Cano, D. Peterse, A. Ghalali, E. Madrian, W. A. Verri, N. Andrews, C. J. Woolf, M. S. Rogers, Pain 2020, 161, 1321–1331.

35. E. O. Adewuyi, D. Mehta, International Endogene Consortium (IEC), 23andMe Research Team, D. R. Nyholt, Hum. Reprod. 2022, 37, 366–383.

36. A. L. Shafrir, M. C. Palmor, J. Fourquet, A. D. DiVasta, L. V. Farland, A. F. Vitonis, H. R. Harris, M. R. Laufer, D. W. Cramer, K. L. Terry, S. A. Missmer, Am. J. Reprod. Immunol. 2021, e13404.

37. N. Shigesi, M. Kvaskoff, S. Kirtley, Q. Feng, H. Fang, J. C. Knight, S. A. Missmer, N. Rahmioglu, K. T. Zondervan, C. M. Becker, Hum. Reprod. Update 2019, 25, 486–503.

38. A. Garitazelaia, A. Rueda-Martínez, R. Arauzo, J. de Miguel, A. Cilleros-Portet, S. Marí, J. R. Bilbao, N. Fernandez-Jimenez, I. García-Santisteban, Life (Basel) 2021, 11, 24.

39. E. Yoshii, H. Yamana, S. Ono, H. Matsui, H. Yasunaga, Am. J. Reprod. Immunol. 2021, 86, e13486.

40. F. Yang, Y. Wu, R. Hockey, J. Doust, G. D. Mishra, G. W. Montgomery, S. Mortlock, medRxiv 2022, doi:10.1101/2022.10.20.22281201.

41. S. Mortlock, R. I. Corona, P. F. Kho, P. Pharoah, J.-H. Seo, M. L. Freedman, S. A. Gayther, M. T. Siedhoff, P. A. W. Rogers, R. Leuchter, C. S. Walsh, I. Cass, B. Y. Karlan, B. J. Rimel, Ovarian Cancer Association Consortium, International Endometriosis Genetics Consortium, G. W. Montgomery, K. Lawrenson, S. P. Kar, Cell Rep. Med. 2022, 3, 100542.

42. S. A. Treloar, D. T. O’Connor, V. M. O’Connor, N. G. Martin, Fertil. Steril. 1999, 71, 701–710.

43. R. Saha, H. J. Pettersson, P. Svedberg, M. Olovsson, A. Bergqvist, L. Marions, P. Tornvall, R. Kuja-Halkola, Fertil. Steril. 2015, 104, 947–952.

44. D. R. Nyholt, S.-K. Low, C. A. Anderson, J. N. Painter, S. Uno, A. P. Morris, S. MacGregor, S. D. Gordon, A. K. Henders, N. G. Martin, J. Attia, E. G. Holliday, M. McEvoy, R. J. Scott, S. H. Kennedy, S. A. Treloar, S. A. Missmer, S. Adachi, K. Tanaka, Y. Nakamura, G. W. Montgomery, Nat. Genet. 2012, 44, 1355–1359.

45. L. Pagliardini, D. Gentilini, P. Vigano’, P. Panina-Bordignon, M. Busacca, M. Candiani, A. M. Di Blasio, J. Med. Genet. 2013, 50, 43–46.

46. T. Kato, K. Yasuda, K. Matsushita, K. J. Ishii, S. Hirota, T. Yoshimoto, H. Shibahara, Front. Immunol. 2019, 10, 2021.

47. B. Peng, F. T. Alotaibi, S. Sediqi, M. A. Bedaiwy, P. J. Yong, Hum. Reprod. 2020, 35, 901–912.

48. H. Malvezzi, C. Hernandes, C. A. Piccinato, S. Podgaec, Reproduction 2019, 158, 1–12.

49. F. El Idrissi, M. Fruchart, K. Belarbi, A. Lamer, E. Dubois-Deruy, M. Lemdani, A. L. N’Guessan, B. C. Guinhouya, D. Zitouni, Front Endocrinol (Lausanne) 2022, 13, 869053.

50. A. Cui, T. Huang, S. Li, A. Ma, J. L. Pérez, C. Sander, D. B. Keskin, C. J. Wu, E. Fraenkel, N. Hacohen,Nature 2024, 625, 377–384.

51. C. A. Dinarello, Immunol. Rev. 2018, 281, 8–27.

52. J. Vallvé-Juanico, X. Santamaria, K. C. Vo, S. Houshdaran, L. C. Giudice, Fertil. Steril. 2019, 112, 1118–1128.

53. R. Sullender, R. K. Agarwal, M. B. Jacobs, H. Valentine, L. Foster, S. K. Agarwal, J. Minim. Invasive Gynecol. 2023, 30, S107.

54. F. T. Alotaibi, S. Sediqi, C. Klausen, M. A. Bedaiwy, P. J. Yong, F&S Science 2023, 4, 47–55.

55. I. M. McGrath, G. W. Montgomery, S. Mortlock, Hum. Reprod. Update 2023, 29, 655–674.

56. K. J. A. Johnston, M. J. Adams, B. I. Nicholl, J. Ward, R. J. Strawbridge, A. Ferguson, A. M. McIntosh, M. E. S. Bailey, D. J. Smith, PLoS Genet. 2019, 15, e1008164.

57. Y. Han, Q. Jia, P. S. Jahani, B. P. Hurrell, C. Pan, P. Huang, J. Gukasyan, N. C. Woodward, E. Eskin, F. D. Gilliland, O. Akbari, J. A. Hartiala, H. Allayee, Nat. Commun. 2020, 11, 1776.

58. M. Ahmed, V.-P. Mäkinen, A. Mulugeta, J. Shin, T. Boyle, E. Hyppönen, S. H. Lee, Commun. Biol. 2022, 5, 614.

59. K. Ishigaki, S. Sakaue, C. Terao, Y. Luo, K. Sonehara, K. Yamaguchi, T. Amariuta, C. L. Too, V. A. Laufer, I. C. Scott, S. Viatte, M. Takahashi, K. Ohmura, A. Murasawa, M. Hashimoto, H. Ito, M. Hammoudeh, S. A. Emadi, B. K. Masri, H. Halabi, S. Raychaudhuri, Nat. Genet. 2022, 54, 1640–1651.

60. C. Eijsbouts, T. Zheng, N. A. Kennedy, F. Bonfiglio, C. A. Anderson, L. Moutsianas, J. Holliday, J. Shi, S. Shringarpure, 23andMe Research Team, A.-I. Voda, Bellygenes Initiative, G. Farrugia, A. Franke, M. Hübenthal, G. Abecasis, M. Zawistowski, A. H. Skogholt, E. Ness-Jensen, K. Hveem, M. Parkes, Nat. Genet. 2021, 53, 1543–1552.

61. H. Lango Allen, K. Estrada, G. Lettre, S. I. Berndt, M. N. Weedon, F. Rivadeneira, C. J. Willer, A. U. Jackson, S. Vedantam, S. Raychaudhuri, T. Ferreira, A. R. Wood, R. J. Weyant, A. V. Segrè, E. K. Speliotes, E. Wheeler, N. Soranzo, J.-H. Park, J. Yang, D. Gudbjartsson, et al., Nature 2010, 467, 832–838.

62. C. S. Gallagher, N. Mäkinen, H. R. Harris, N. Rahmioglu, O. Uimari, J. P. Cook, N. Shigesi, T. Ferreira, D. R. Velez-Edwards, T. L. Edwards, S. Mortlock, Z. Ruhioglu, F. Day, C. M. Becker, V. Karhunen, H. Martikainen, M. R. Järvelin, R. M. Cantor, P. M. Ridker, K. L. Terry, C. C. Morton, dNat. Commun. 2019, 10, 4857.

63. E. O. Dareng, S. G. Coetzee, J. P. Tyrer, P.-C. Peng, W. Rosenow, S. Chen, B. D. Davis, F. S. Dezem, J.-H. Seo, R. Nameki, A. L. Reyes, K. K. H. Aben, H. Anton-Culver, N. N. Antonenkova, G. Aravantinos, E. V. Bandera, L. E. Beane Freeman, M. W. Beckmann, A. Beeghly-Fadiel, J. Benitez, S. A. Gayther, Am. J. Hum. Genet. 2024, 111, 1061–1083.

64. K. Watanabe, S. Stringer, O. Frei, M. Umićević Mirkov, C. de Leeuw, T. J. C. Polderman, S. van der Sluis, O. A. Andreassen, B. M. Neale, D. Posthuma, Nat. Genet. 2019, 51, 1339–1348.

65. C. A. de Leeuw, J. M. Mooij, T. Heskes, D. Posthuma, PLoS Comput. Biol. 2015, 11, e1004219.

66. K. Watanabe, E. Taskesen, A. van Bochoven, D. Posthuma, Nat. Commun. 2017, 8, 1826.

67. S. Yazar, J. Alquicira-Hernandez, K. Wing, A. Senabouth, M. G. Gordon, S. Andersen, Q. Lu, A. Rowson, T. R. P. Taylor, L. Clarke, K. Maccora, C. Chen, A. L. Cook, C. J. Ye, K. A. Fairfax, A. W. Hewitt, J. E. Powell, Science 2022, 376, eabf3041.

68. C. E. Filby, K. A. Wyatt, S. Mortlock, F. L. Cousins, B. McKinnon, K. E. Tyson, G. W. Montgomery, C. E. Gargett, J. Pers. Med. 2021, 11, doi:10.3390/jpm11121314.

69. M. Y. Turco, L. Gardner, J. Hughes, T. Cindrova-Davies, M. J. Gomez, L. Farrell, M. Hollinshead, S. G. E. Marsh, J. J. Brosens, H. O. Critchley, B. D. Simons, M. Hemberger, B.-K. Koo, A. Moffett, G. J. Burton, Nat. Cell Biol. 2017, 19, 568–577.

70. B. D. McKinnon, S. W. Lukowski, S. Mortlock, J. Crawford, S. Atluri, S. Subramaniam, R. L. Johnston, K. Nirgianakis, K. Tanaka, A. Amoako, M. D. Mueller, G. W. Montgomery, Commun. Biol. 2022, 5, 600.

71. C. E. Gargett, K. E. Schwab, R. M. Zillwood, H. P. T. Nguyen, D. Wu, Biol. Reprod. 2009, 80, 1136–1145.

72. S. Gurung, J. A. Werkmeister, C. E. Gargett, Sci. Rep. 2015, 5, 15042.

73. R. González-Cano, M. Merlos, J. M. Baeyens, C. M. Cendán, Anesthesiology 2013, 118, 691–700.

74. R. Gonzalez-Cano, B. Boivin, D. Bullock, L. Cornelissen, N. Andrews, M. Costigan, Front. Pharmacol. 2018, 9, 433.

75. J. M. A. Laird, L. Martinez-Caro, E. Garcia-Nicas, F. Cervero, Pain 2001, 92, 335–342.

76. V. Fattori, F. A. Pinho-Ribeiro, S. M. Borghi, J. C. Alves-Filho, T. M. Cunha, F. Q. Cunha, R. Casagrande, W. A. Verri, Inflamm. Res. 2015, 64, 993–1003.

