## Supplemental figures for "M2 Macrophages are Major Mediators of Germline Risk of Endometriosis and Explain Pleiotropy with Comorbid Traits"

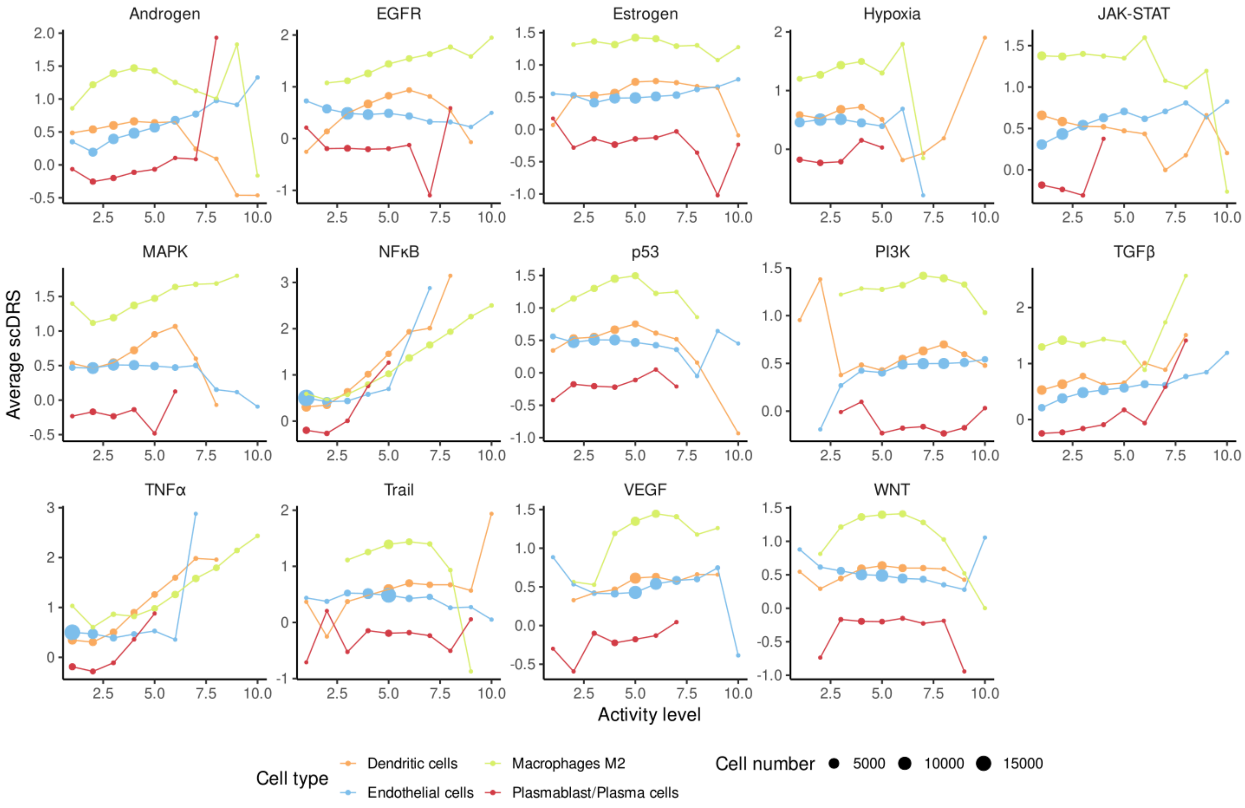


**Supplementary figure 1. scDRS heterogeneity relative to pathway activity.** While most cells have low activity levels for all pathways, matching the expected for single cell data, M2 macrophages cluster around intermediate levels of NFκB and TNF-α.

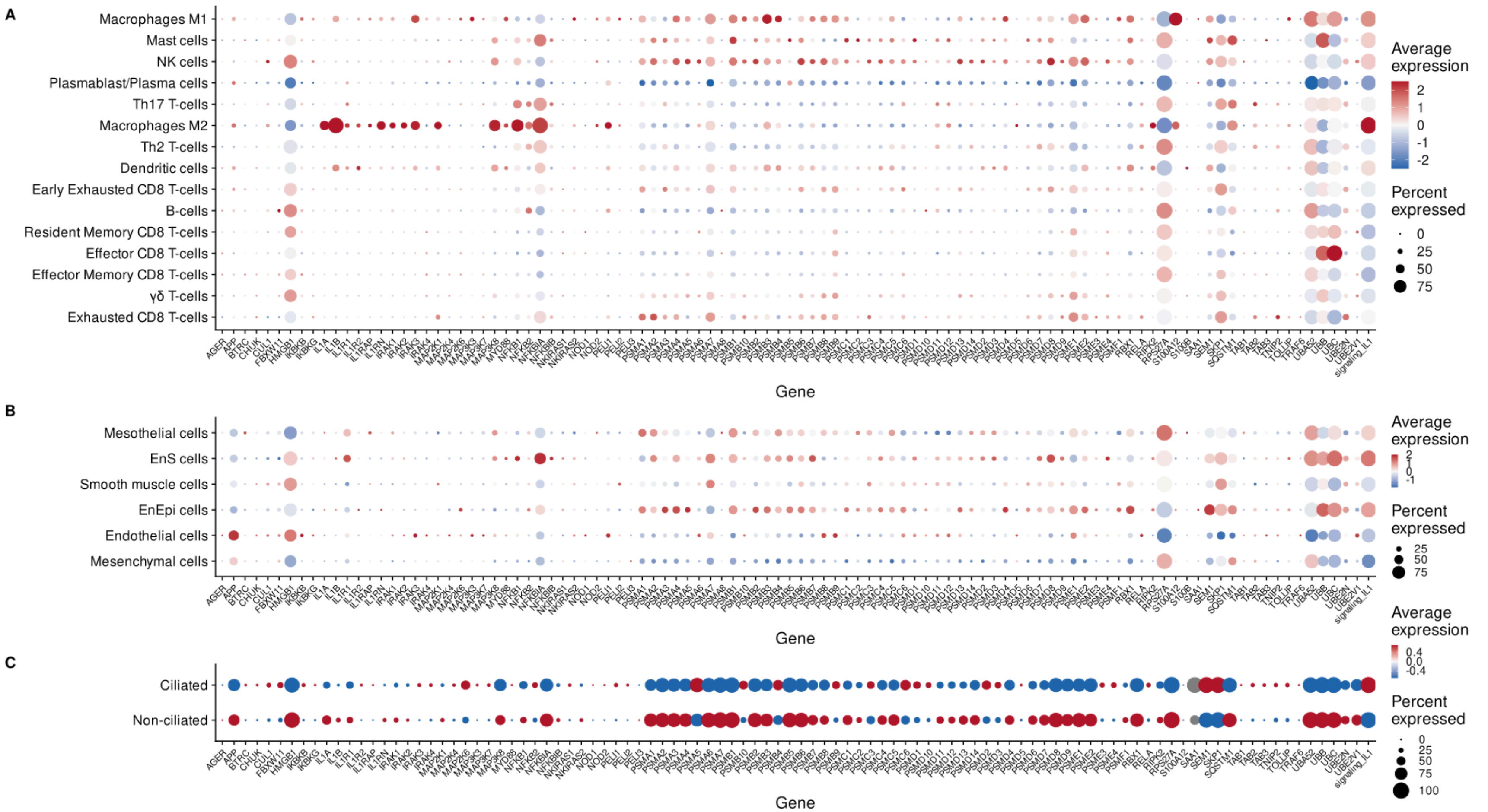


**Supplementary figure 2. Expression of the IL1 signature and associated genes per cell cluster. A**) immune subsets from patient single cell data. **B**) non-immune subsets from patient single cell data. **C**) organoid single cell data

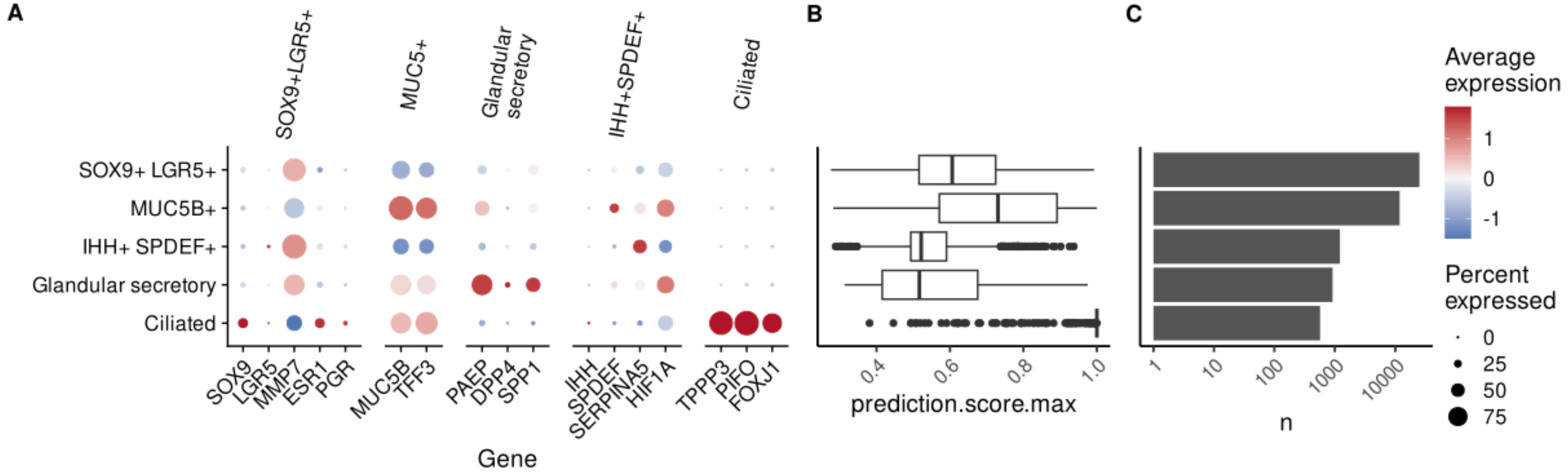


**Supplementary figure 3. Label transfer results for the endometrial epithelial subclusters. A**) Expression of markers from the endometrial epithelial subclusters. **B**) Label assignment score. **C**) Number of cells in each cluster
